# Chromosome-scale genome assembly and annotation of the two-spotted cricket *Gryllus bimaculatus* (Orthoptera: Gryllidae)

**DOI:** 10.1101/2025.10.31.685973

**Authors:** Kosuke Kataoka, Ryuto Sanno, Tomasz Gaczorek, Upendra Raj Bhattarai, Yuki Ito, Shintaro Inoue, Kei Yura, Toru Asahi, Guillem Ylla, Taro Mito, Cassandra G. Extavour

## Abstract

The two-spotted cricket, *Gryllus bimaculatus*, is a key hemimetabolous model organism for developmental biology, neuroscience, and regeneration. The existing reference genome is, however, highly fragmented into 47,877 scaffolds, hampering chromosome-scale analyses for these fields. Here, we report a high-quality, chromosome-scale genome assembly for the white-eyed mutant strain of this cricket, generated using a combination of Nanopore and PacBio HiFi long reads, integrated with Hi-C data. The final 1.62 Gbp assembly achieves a scaffold N50 of 107.4 Mbp, a significant improvement in contiguity over the previous 6.3 Mbp N50. We anchored 94.45% of the assembly into 15 pseudomolecules, consistent with the known karyotype (n = 15). The genome completeness (BUSCO v6.0.0 insecta_odb12) reached 98.1%. We also updated the annotation, identifying 14,964 protein-coding genes. This gene set shows markedly improved completeness (BUSCO v6.0.0 insecta_odb12: 95.7%) compared with the previous annotation (81.2%) and successfully recovers all nine essential neuropeptide genes previously reported as missing from the draft assembly. This chromosome-scale genomic resource provides an essential foundation for comparative and functional genomics in *G. bimaculatus*.

## Introduction

The two-spotted cricket, *Gryllus bimaculatus*, is a model organism for hemimetabolous insects (Horch et al., 2017) (Fig. 1). Unlike holometabolous insects such as the fruit fly (*Drosophila melanogaster*) or the red flour beetle (*Tribolium castaneum*), hemimetabolous insects undergo direct development, where nymphs hatch and grow through successive molts to become adults without larval or pupal stages. This ancestral developmental mode is crucially important for understanding insect evolution. Due to its ease of rearing, short generation time, and the applicability of efficient RNA interference (RNAi) (Miyawaki et al., 2004) and genome-editing techniques such as CRISPR/Cas9 (Matsuoka et al., 2025), *G. bimaculatus* has been established as a powerful experimental model in a wide range of fields, including evolutionary developmental biology (Donoughe & Extavour, 2016), regeneration biology (Nakamura et al., 2008), neuroscience (Matsumoto et al., 2018), and ethology (Abe et al., 2021; Kuriwada, 2022). It is also gaining significant global attention as a novel, sustainable food source due to its high protein content and efficient rearing (Kataoka et al., 2020, 2022; Mito et al., 2022).

**Figure 1.**
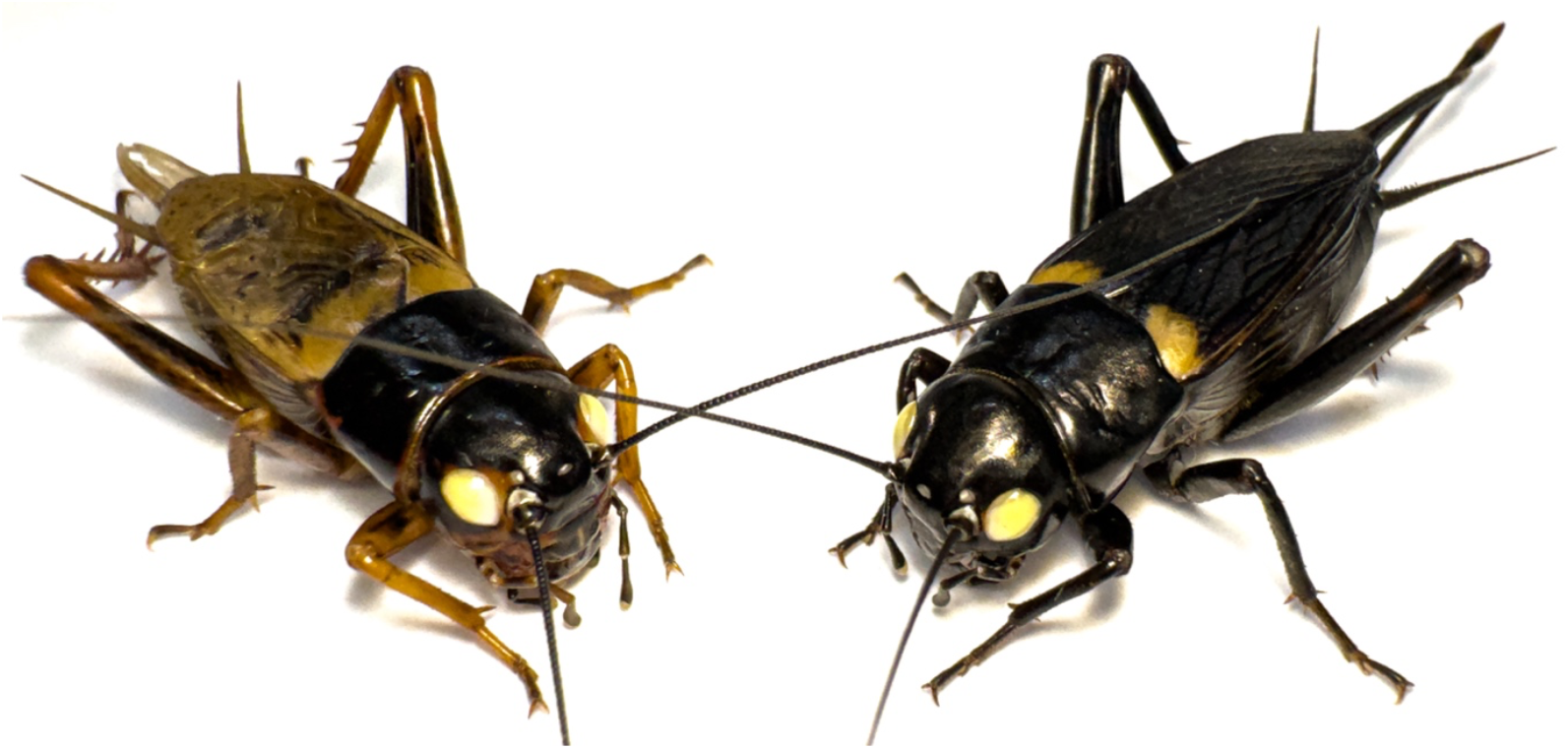
Adult male (left) and female (right) of the white-eyed mutant strain *G. bimaculatus*.

In the first report of a genomic resource for *G. bimaculatus*, the white-eyed mutant strain, which is standardly used in many functional studies, was sequenced (Ylla et al., 2021). The assembly (GenBank accession: GCA_017312745.1) was a useful resource with a genome size of approximately 1.66 Gb and a scaffold N50 of 6.3 Mb, contributing to analyses of gene family evolution and DNA methylation. However, this previous assembly was fragmented into 47,877 scaffolds and was not assembled to chromosome scale. While chromosome-scale genomes have recently been reported for other related cricket species, such as *Gryllus assimilis* (Ito et al., 2025) and *Acheta domesticus* (Dossey et al., 2023), such a resource has remained unavailable for *G. bimaculatus*, which serves as the primary model for functional, developmental, and neurobiological studies. The lack of a chromosome-scale assembly for this specific species has been a significant limitation for large-scale comparative genomic analyses based on synteny (conservation of gene order), understanding the structural arrangement of transposons and repetitive sequences on chromosomes, and for quantitative trait locus mapping and the accurate identification of genome-editing off-target sites.

In this study, to fill the gap, we constructed a high-quality, chromosome-scale genome assembly using the same, white-eyed mutant strain, used in the previous study. By combining Nanopore and PacBio HiFi long-read sequencing methods, and Hi-C chromatin conformation capture technology, we anchored 94.45% of the entire genome into 15 pseudomolecules, corresponding to the *G. bimaculatus* karyotype (n = 14 autosomes + X) (Yoshimura et al., 2006). This assembly achieves a contig N50 of 4.58 Mb and a scaffold N50 of 107.39 Mb, representing a significant improvement in contiguity. Furthermore, we report an updated gene annotation comprising 14,964 protein-coding genes. This chromosome-scale genome resource will strengthen the foundation for all genomic research using *G. bimaculatus* and accelerate new insights into the biology of hemimetabolous insects.

## Materials and Methods

### Animals

A white-eyed mutant strain of *G. bimaculatus* (Mito & Noji, 2008) was housed in plastic cases at 30 °C ± 1 °C and 30%–40% relative humidity under a 10 h light and 14 h dark photoperiod. They were nourished with an artificial fish food (4971618–011312, Kyorin, Japan).

### Library preparation and sequencing

A single alive *G. bimaculatus* individual (Fig. 1) was used for genomic DNA extraction. Total genomic DNA was extracted from the head and hind legs of a male *G. bimaculatus* using NucleoBond® HMW DNA (Macherey-Nagel, Germany) according to the manufacturer’s instructions. The resulting genomic DNA was size-selected using a Short Read Eliminator Kit (PacBio, CA, USA). DNA purity and concentrations were measured by spectrometry using NanoPhotometer NP80-TOUCH (Implen, Germany) and fluorometry using Qubit 4 (Thermo Fisher Scientific, MA, USA).

For long-read sequencing, Oxford Nanopore Technologies (ONT) libraries were constructed using the Ligation Sequencing Kit V14 and sequenced on the PromethION 2 Solo platform (Oxford Nanopore Technologies, UK) with a Flow Cell R10.4.1. Base-calling was performed using Dorado v0.3.0 (model: dna_r10.4.1_e8.2_400bps_sup@v4.2.0). The resulting raw reads were adapter-trimmed using Porechop_ABI v0.5.0 (Bonenfant et al., 2023). Additionally, a SMRTbell library was prepared and sequenced on a PacBio Sequel IIe system.

For chromosome-scale scaffolding, two separate Hi-C libraries were prepared. The first library was prepared from the hind legs of a single male *G. bimaculatus* using the Dovetail™ Omni-C™ Kit (Dovetail Genomics, CA, USA) following the manufacturer’s instructions. The second Hi-C library was generated from the thorax and legs of an adult male using the Proximo Hi-C (Animal) kit (KT2045) (Phase Genomics, Seattle, US), following the manufacturer’s instructions. Both libraries were sequenced on the Illumina NovaSeq 6000 platform, and the sequencing data were combined for downstream scaffolding analysis.

### Genome *de novo* assembly

The initial draft genome was assembled by combining the filtered ONT long reads and the PacBio HiFi reads using Flye v2.9.5 (Kolmogorov et al., 2019). To correct errors in the resulting contigs, we retrieved publicly available Illumina short-read data (DDBJ Sequence Read Archive [DRA] accessions DRR272308–DRR272313), which were originally sequenced on an Illumina HiSeq 2000. These reads were downsampled to an approximate 93.9x coverage and used for polishing. The assembly underwent two rounds of error correction using POLCA v4.1.0 (Zimin & Salzberg, 2020) with default settings.

Potential contamination in the assembly was removed using BlobToolKit v1.1.1 (Challis et al., 2020), which analyzes unexpected coverage, GC content, or similarity to bacterial and other contaminant sequences. Sequence coverage was determined by mapping Illumina reads with bwa v0.7.17-r1188. Similarity analysis was performed using BLASTn v2.13.0+ against NCBI NT database v5 (options: - task megablast culling_limit 10 -evalue 1e-25 -outfmt ‘6 qseqid staxids bitscore std sscinames sskingdoms stitle’). Mitochondrial genomes were also identified through gene prediction using the MITOS2 webserver and subsequently removed.

The resulting contigs were then corrected for misjoins, ordered, oriented, and anchored into a chromosome-scale assembly using Omni-C™ data with Juicer v1.9.9 (Durand et al., 2016) and 3D-DNA v180419 (Dudchenko et al., 2017). Candidate assembly was reviewed with Juicebox Assembly Tools v1.9.9 (Durand et al., 2016) for quality control and interactive corrections. The contact map was visualized using Juicebox. The completeness of the final genome assembly was assessed using BUSCO v6.0.0 (Tegenfeldt et al., 2025) against the insecta_odb12 lineage datasets.

### Prediction of repeat regions

In *de novo* repeat prediction, RepeatModeler v2.0.6 (Flynn et al., 2020) was first used for *de novo* repeat identification. Because standard libraries are insufficient for effective masking in *Gryllus* genomes (Szrajer et al., 2024), this library was supplemented with the custom repeat library previously generated (Ylla et al., 2021). The combined repeat library was then used to identify and softmask repetitive elements in the *G. bimaculatus* genome using RepeatMasker v4.2.1 (Smit et al., 2015).

### Structural gene annotation

Structural annotation for protein-coding genes was performed on the softmasked genome using *ab initio* prediction and RNA-seq-based prediction. Both methods used publicly available RNA-seq data (Sequence Read Archive [SRA] accessions: SRR10619411, SRR10619415, SRR10619417, SRR10619418, SRR10619421, SRR10619423, SRR10619425, SRR10619429, SRR10619431, SRR10619432, SRR10619434, SRR10619437, SRR10619439, SRR10619440, SRR14026720– SRR14026726) as input.

To remove noisy RNA-seq reads potentially arising from erroneous transcription and splicing, *de novo* transcriptome assembly was first performed using Trinity v2.15.1 (Haas et al., 2013) to generate contigs. The original RNA-seq reads were then mapped back to these contigs using HISAT2 v2.2.1 (Kim et al., 2019) with default parameters, allowing filtration of reads that did not map correctly in the proper orientation as paired-end reads. After removing these noisy reads, the remaining reads were subsequently used for gene predictions.

The *ab initio* prediction was carried out using BRAKER v3.0.8 (Brůna et al., 2021; Gabriel et al., 2024; Hoff et al., 2016, 2019; Stanke et al., 2008, 2006), incorporating protein data from the OrthoDB 12 arthropods dataset (retrieved from https://bioinf.uni-greifswald.de/bioinf/partitioned_odb12/) and the mapping data of the filtered RNA-seq reads. This BRAKER prediction served as the foundation for our gene set. To complement this base set, StringTie2 v2.2.1 (Kovaka et al., 2019) was used for RNA-seq-based prediction. These combined predictions (BRAKER and StringTie2) were merged and duplicate genes were discarded using GffCompare v0.12.6 (Pertea & Pertea, 2020) to form a final, comprehensive consensus gene set.

### Functional gene annotation

Gene functional annotation was conducted using eggNOG-mapper online (http://eggnog-mapper.embl.de/) (Cantalapiedra et al., 2021) and BLASTp-based methods. For the BLASTp-based annotation, we used databases including *Homo sapiens, Mus musculus, Caenorhabditis elegans, D. melanogaster*, and UniProt Swiss-Prot to identify the best hits for annotation (E-value < 1.0 × 10^−10^).

The completeness of the final predicted gene models was assessed using BUSCO v6.0.0 (Tegenfeldt et al., 2025) against the insecta_odb12 lineage dataset. For this analysis, the longest isoform for each gene was first extracted from the annotation file using AGAT v0.9.1 (Dainat et al., 2022).

### Validation of neuropeptide gene loci

To assess the completeness of our assembly regarding functionally important gene families, we specifically investigated the neuropeptide gene loci previously reported (Mochizuki et al., 2023). The neuropeptide cDNA sequences listed in the study were retrieved. These sequences were mapped against our final chromosome-scale genome assembly using Exonerate v2.4.0 (Slater & Birney, 2005) with the est2genome model to accurately determine exon-intron boundaries. All resulting alignments were manually inspected to validate the gene structures.

## Results and Discussion

### Genome sequencing

To construct the genome assembly, we generated three types of sequencing data (Table 1). First, we obtained 45.00 Gbp of ONT sequencing data; the average and N50 read lengths were 13.13 Kbp and 24.23 Kbp, respectively. Second, we generated 13.44 Gbp of PacBio HiFi data, with an average read length of 13.61 Kbp and an N50 of 14.17 Kbp. Finally, for chromosome-scale scaffolding, 243.22 Gbp of Hi-C raw data was generated from the Illumina platform.

**Table 1.**
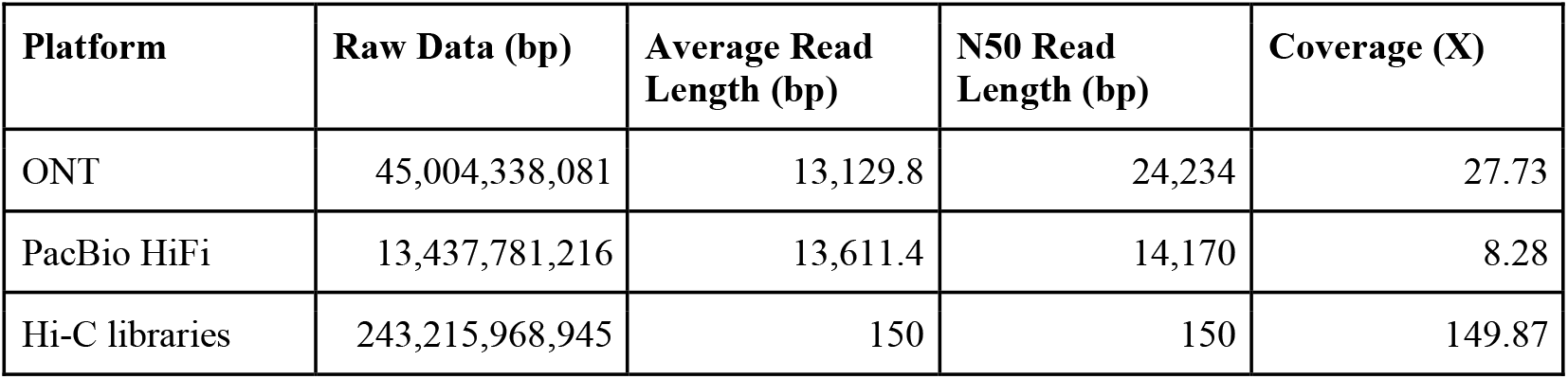
Statistics for the DNA-seq data of the *G. bimaculatus* genome.

| Platform | Raw Data (bp) | Average Read Length (bp) | N50 Read Length (bp) | Coverage (X) |
| --- | --- | --- | --- | --- |
| ONT | 45,004,338,081 | 13,129.8 | 24,234 | 27.73 |
| PacBio HiFi | 13,437,781,216 | 13,611.4 | 14,170 | 8.28 |
| Hi-C libraries | 243,215,968,945 | 150 | 150 | 149.87 |

### Genome *de novo* assembly statistics

The hybrid assembly strategy combining ONT and HiFi reads, followed by two rounds of Illumina-based polishing, yielded a 1.63 Gbp draft genome. This polished assembly comprised 3,789 contigs and achieved a contig N50 of 4.58 Mbp.

Following Hi-C-based scaffolding, the final assembly resulted in a genome of 1.62 Gbp in size, visualized by a snail plot (Fig. 2A). This assembly consists of 196 scaffolds with a scaffold N50 length of 107 Mbp (Table 2). This represents a substantial improvement in contiguity compared to the previous *G. bimaculatus* assembly (Ylla et al., 2021), which had a scaffold N50 of 6.3 Mbp and was fragmented into 47,877 scaffolds (Fig. 2B). To assess genomic completeness, we performed a BUSCO v6.0.0 analysis using the insecta_odb12 dataset. Our assembly achieved a completeness score of 98.1% (C:98.1%[S:96.0%,D:2.2%],F:0.4%,M:1.5%), demonstrating a higher level of completeness compared to the 96.0% (C:96.0%[S:94.3%,D:1.7%],F:1.4%,M:2.6%) of the previous genome (Table 3).

**Table 2.** Summary statistics for the chromosome-scale assembly of *G. bimaculatus*.

| Features | Statistic |
| --- | --- |
| Number of chromosomes | 15 |
| Number of scaffolds | 196 |
| Scaffold N50 (bp) | 107,386,002 |
| GC content (%) | 40.36 |
| Max scaffold size (bp) | 254,522,992 |
| Total size (bp) | 1,622,898,960 |

**Table 3.** BUSCO completeness assessment of the *G. bimaculatus* genome assembly.

| BUSCO v6.0.0 | This Study | Ylla et al. 2021 |
| --- | --- | --- |
| Complete BUSCOs | 98.1% | 96.0% |
| Single-copy complete BUSCOs | 96.0% | 94.3% |
| Duplicated complete BUSCOs | 2.2% | 1.7% |
| Fragmented BUSCOs | 0.4% | 1.4% |
| Missing BUSCOs | 1.5% | 2.6% |

**Figure 2.**
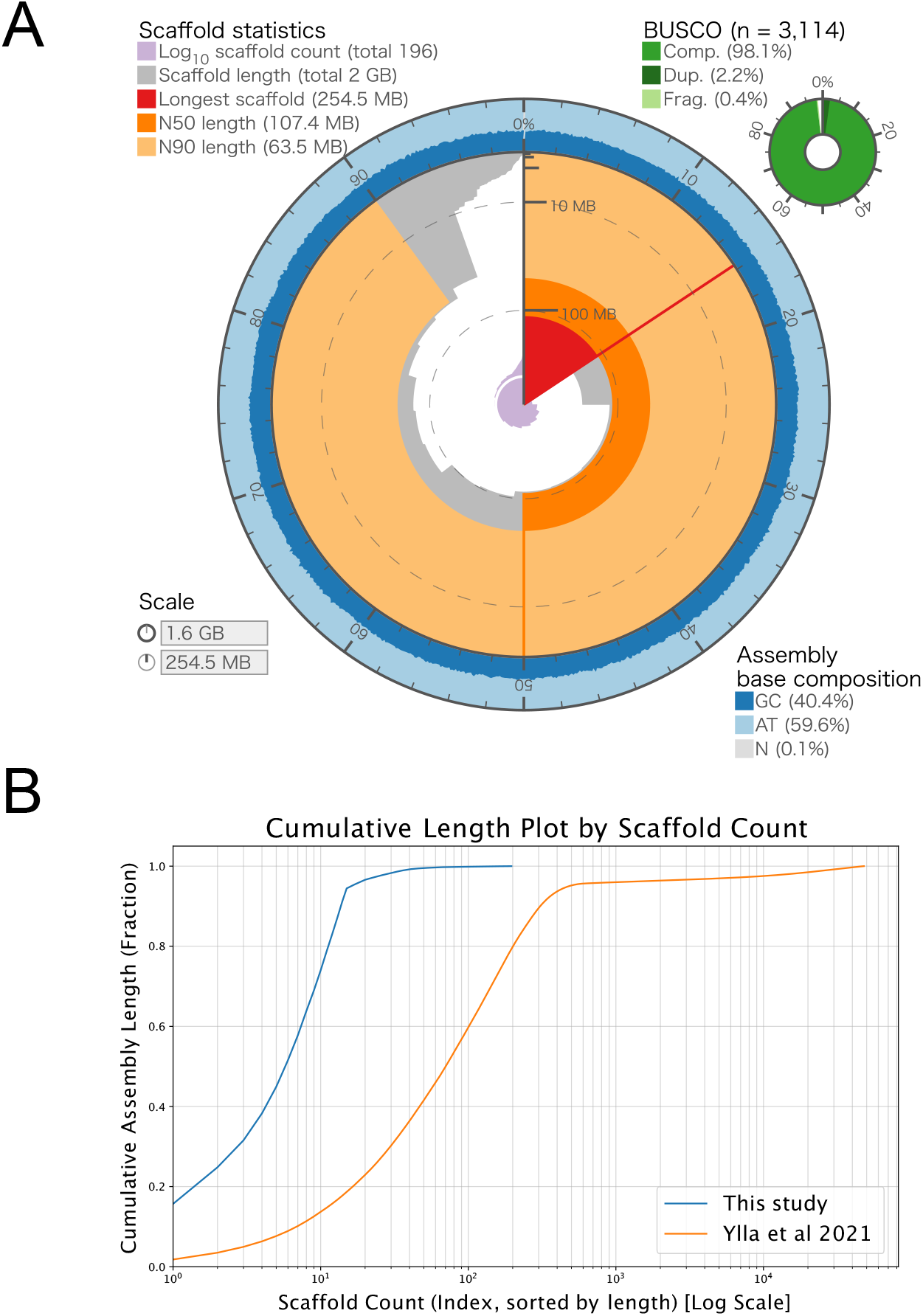
Assembly statistics and contiguity of the *Gryllus bimaculatus* (white-eyed strain) chromosome-scale genome. **(A)** Snail plot visualizing the key statistics of the final assembly. The cumulative assembly length (1.62 Gbp) is plotted in light orange, with the longest scaffold (254.5 Mbp) shown in red. The N50 length (107.4 Mbp) is indicated by the orange arc. The inner purple spiral plots the cumulative number of scaffolds (196 total) on a log scale, with white scale lines drawn at successive orders of magnitude from 10 scaffolds onwards. The circumferential axis indicates the base composition of the assembly (GC: 40.4%, AT: 59.6%, N: 0.1%). The donut chart (top right) displays the BUSCO completeness (insecta_odb12), showing 98.1% total complete (“Comp.”) genes (light green), which includes 96.0% single-copy and 2.2% duplicated (“Dup.”) genes (dark green). An additional 0.4% were fragmented (“Frag.”) genes (pale green). An interactive version is available at https://kataokaklab.github.io/snailplot-assembly-stats/. **(B)** Cumulative length plot comparing the contiguity of this assembly (blue line) with the previous draft assembly (Ylla et al., 2021) (orange line). The y-axis shows the percentage of the total genome length covered, and the x-axis (log scale) shows the number of scaffolds.

A total of 15 pseudochromosomes were constructed and accounted for 94.45% of the total genome length (Table 4). This number (n = 15) is in agreement with the established karyotype for *G. bimaculatus* (n = 14 autosomes + X), which was previously determined by cytogenetic analysis (Yoshimura et al., 2006) and is also consistent with that of the recently reported species *G. assimilis* (Ito et al., 2025), which is closely related to *G. bimaculatus*. The X chromosome was identified as the longest, accounting for 15.68% of the genome, which is in accordance with the karyotype of this species (Yoshimura et al., 2006). This was further validated by the observation that the X chromosome displayed half of the read coverage compared to the autosomal chromosomes, calculated using a genomic short-read library (DRR272308) from a single hemizygous male (Fig. S1).

**Table 4.**
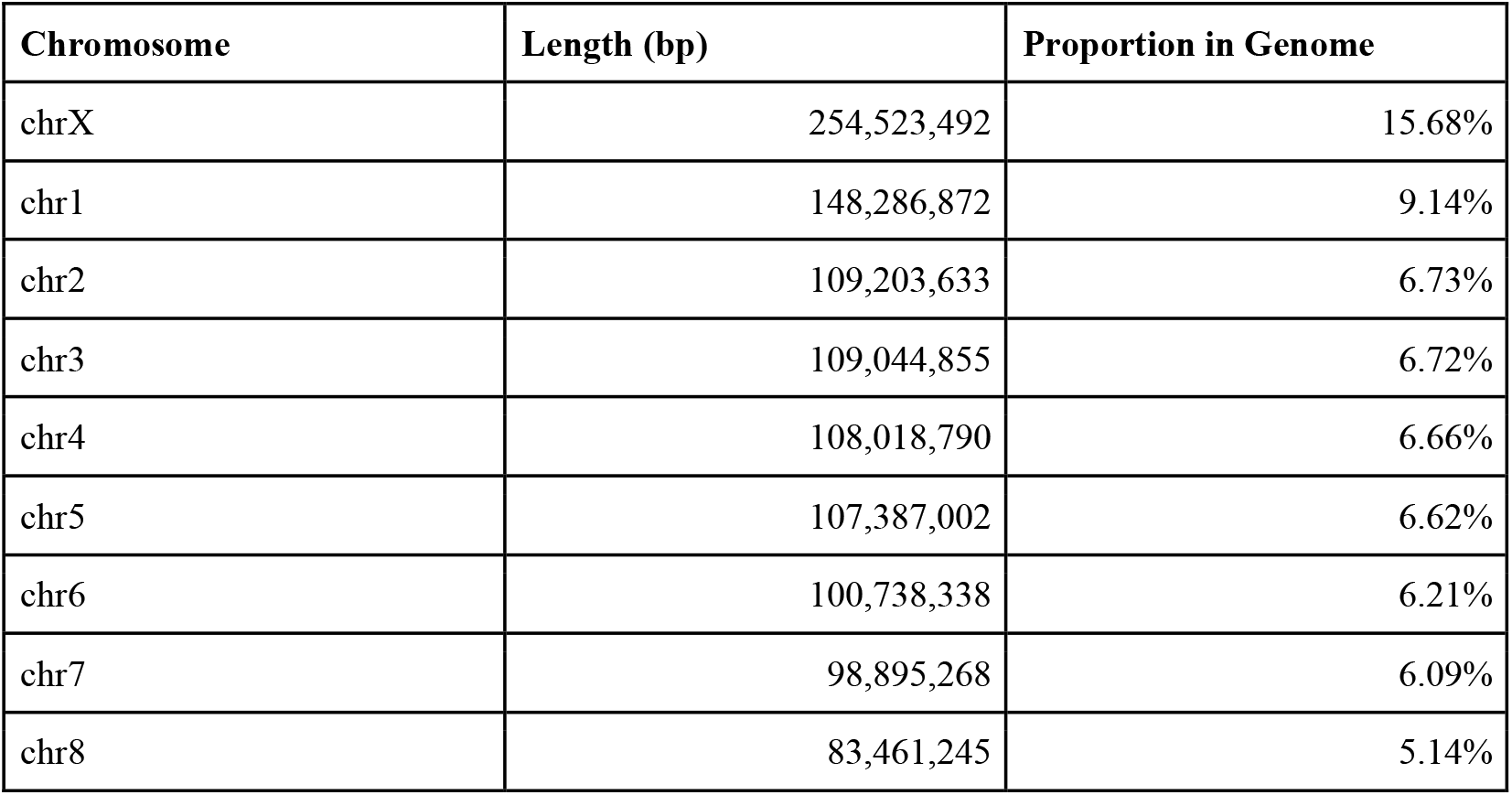
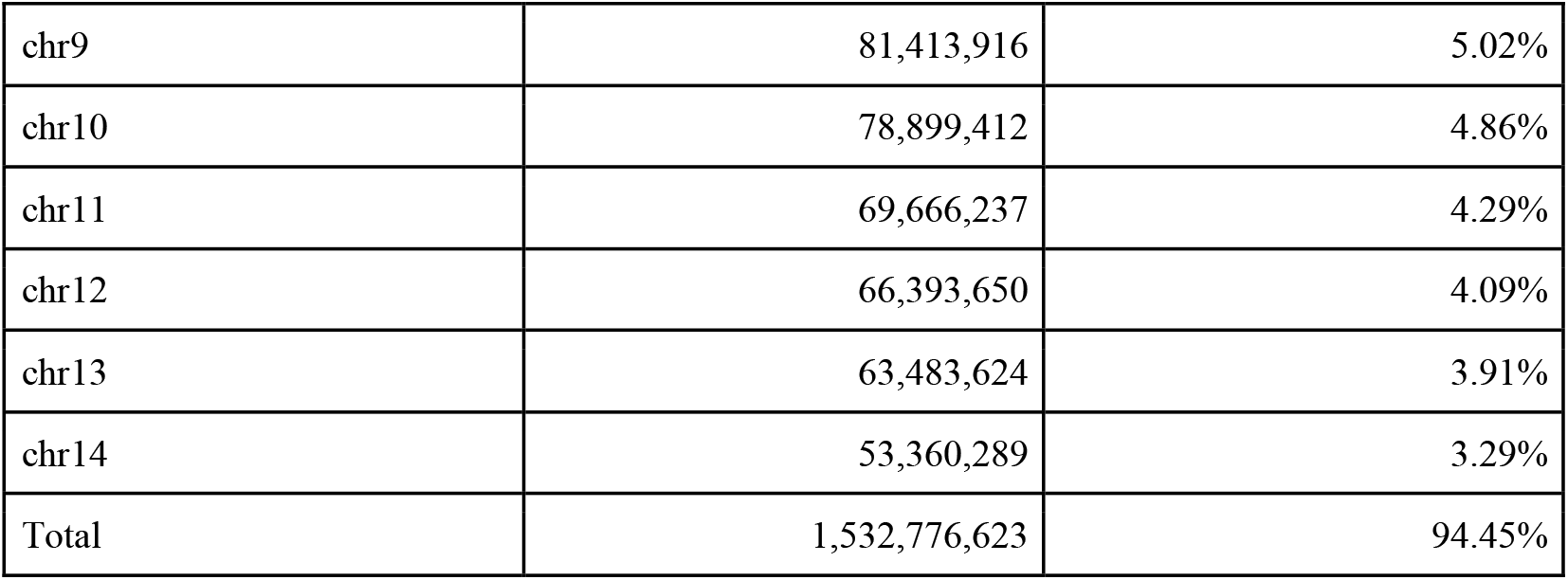
Statistics of the assembled pseudochromosomes in *G. bimaculatus*.

### Gene annotation

The final consensus gene set, derived from merging BRAKER *ab initio* predictions and StringTie2 RNA-seq-based predictions, comprised 14,964 protein-coding genes (Table 5). This total is less than the 17,871 genes reported previously (Ylla et al. 2021), likely because the improved scaffold contiguity of our assembly allows for the correct assembly of genes previously split across multiple scaffolds. Of the 14,964 predicted genes, functional annotation was assigned using eggNOG-mapper and BLASTp (E-value < 1.0 ×10^−10^ against several model organisms’ annotation datasets. eggNOG-mapper annotated 71.96% of the genes. The BLASTp searches (E-value < 1.0 × 10^−10^) against databases including *H. sapiens, M. musculus, C. elegans, D. melanogaster*, and UniProt Swiss-Prot yielded hit rates ranging from 44.84% to 71.89% (Table 6).

**Table 5.**
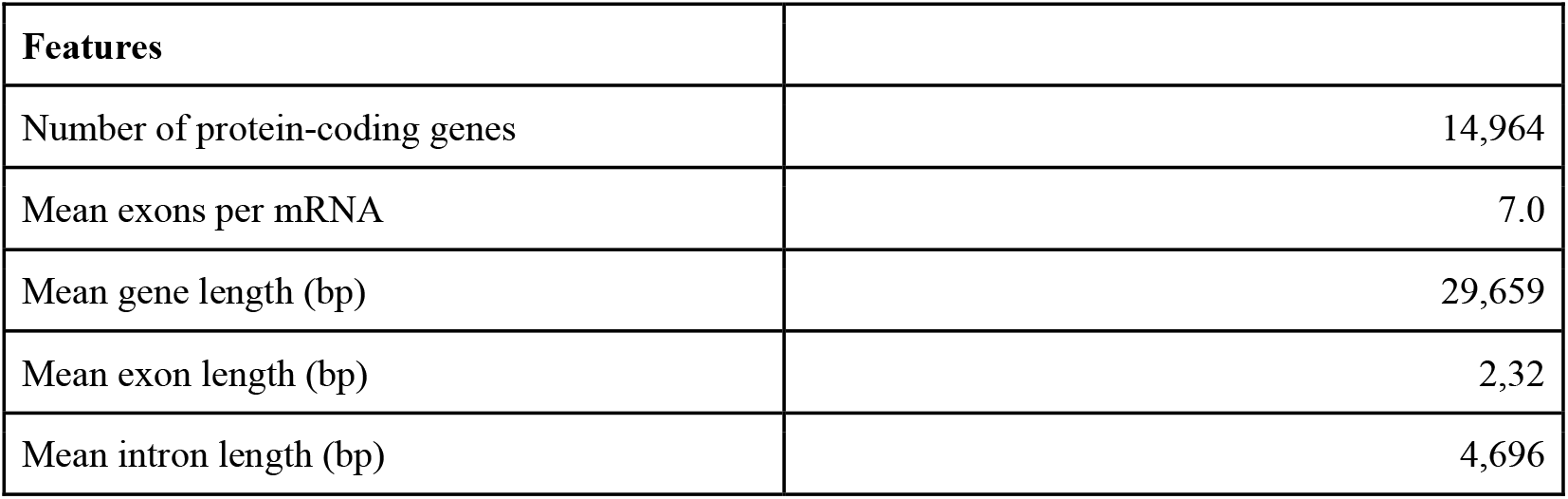
Summary statistics for gene prediction in *G. bimaculatus*.

| Features |  |
| --- | --- |
| Number of protein-coding genes | 14,964 |
| Mean exons per mRNA | 7.0 |
| Mean gene length (bp) | 29,659 |
| Mean exon length (bp) | 2,32 |
| Mean intron length (bp) | 4,696 |

**Table 6.** Structural statistics of the *G. bimaculatus* gene set.

| Annotation Sources |  |
| --- | --- |
| *Homo sapiens* (GRCh38) | 44.84% |
| *Mus musculus* (GRCm39) | 71.89% |
| *Caenorhabditis elegans* (WBcel235) | 55.18% |
| *Drosophila melanogaster* (BDGP6.32) | 67.16% |
| Uniprot/Swissprot (release: 2020_06) | 66.74% |
| eggNOG-mapper | 71.96% |

The completeness of this annotation set was validated using BUSCO v6.0.0. The analysis identified 95.7% of the expected complete insecta BUSCOs (C:95.7%[S:93.6%,D:2.0%],F:1.5%,M:2.9%) (Table 7). These results collectively indicate a high-quality gene set for this species.

**Table 7.** Functional annotation summary for the *G. bimaculatus* gene set.

| BUSCO v6.0.0 | This study | Ylla et al. 2021 |
| --- | --- | --- |
| Complete BUSCOs | 95.7% | 81.2% |
| Single-copy complete BUSCOs | 93.6% | 79.9% |
| Duplicated complete BUSCOs | 2.0% | 1.3% |
| Fragmented BUSCOs | 1.5% | 8.9% |
| Missing BUSCOs | 2.9% | 10.0% |

### Recovery of missing neuropeptide genes

A recent comprehensive study (Mochizuki et al., 2023) highlighted significant gaps in the previous draft genome assembly (Ylla et al., 2021). They reported that several crucial neuropeptides genes (e.g., ACP (Adipokinetic hormone/corazonin-related peptide), Allatotropin, Kinin, etc.) were missing from the draft genome and could only be identified within *de novo* transcriptome assemblies, suggesting these loci were absent from the previous reference.

We asked if our new chromosome-scale assembly was more complete in this regard, by mapping the cDNA sequences of these previously missing neuropeptides. We successfully located all nine of these genes encoding neuropeptides (i.e., ACP, Allatostatin CC (Ast CC), Allatotropin, CCHamide-1, CCHamide-2, CRF/DH (Corticotropin releasing factor-like diuretic hormone), Kinin (Leucokinin), Neuropeptide F1a (NPF1a), Neuropeptide F1b (NPF1b)), which are now correctly anchored onto our pseudomolecules. For example, the ACP gene, previously missing, was successfully mapped to Chromosome X, where it spans 11,668 bp and is composed of three exons, revealing its complete exon-intron structure (Fig. S2). This demonstrates that our assembly not only improves contiguity to the chromosome scale but also recovers functionally critical genes that were absent in the previous reference, providing a more complete and reliable resource for functional genomics in *G. bimaculatus*.

## Conclusions

We have generated a high-quality, chromosome-scale genome assembly and updated gene annotation for the key hemimetabolous model organism *G. bimaculatus*. This assembly represents a substantial upgrade to the previous draft sequence (Ylla et al., 2021), increasing the scaffold N50 from 6.3 Mbp to 107.4 Mbp and anchoring 94.45% of the sequence into 15 pseudomolecules, consistent with the known karyotype (Yoshimura et al., 2006).

Crucially, our assembly resolves significant gaps present in the previous version, evidenced by the recovery of nine essential neuropeptide genes previously reported as missing (Mochizuki et al., 2023). This improved completeness is further supported by superior BUSCO scores for both the genome (98.1% vs. 96.0%) and the gene set (95.7% vs. 81.2%). This highly contiguous and complete genome sequence provides an essential new foundation for the *G. bimaculatus* research community, facilitating advanced genetic and genomic analyses, such as synteny comparisons, QTL mapping, and the precise design of genome-editing experiments.

## Data Availability

The scripts used for the analyses in this study are available in GitHub (https://github.com/Kataoka-K-Lab/Gryllus_bimaculatus_genome_gbim_v2.2). All bioinformatics tools used in this study followed their respective manuals and protocols. The software versions, codes, and parameters are provided in the Materials and Methods section. Unless otherwise specified, default parameters were used.

The genomic WGS sequencing data were deposited in the NCBI Sequence Read Archive (SRA) database under the BioProject PRJNA1347939.

The assembled genome and annotation datasets are also available in figshare (https://doi.org/10.6084/m9.figshare.30472754.v1) (Kataoka, 2025).

## Funding

This study was supported by the Cabinet Office, Government of Japan, Cross-ministerial Moonshot Agriculture, Forestry and Fisheries Research and Development Program, “Technologies for Smart Bio-industry and Agriculture” (BRAIN) [JPJ009237] (K.K., S.I., T.A., K.Y., and T.M.), the Strategic Programme Excellence Initiative at the Jagiellonian University – BioS PRA (T.G. and G.Y.), and the National Science Foundation Award [IOS-2220747] (C.G.E.). C.G.E. is an investigator of the Howard Hughes Medical Institute.

## Competing Interests

The authors declare no conflict of interest.

## Supplementary Figures

**Supplementary Figure S1. Genomic read coverage confirms X chromosome hemizygosity**. Mean read coverage depth across all assembled chromosomes derived from the mapping of genomic short reads of a single male *Gryllus bimaculatus* individual (SRA: DRR272308). The X chromosome displays approximately half the average autosomal coverage depth, consistent with the expected pattern of male hemizygosity.

**Supplementary Figure S2. Recovery of the Adipokinetic hormone/corazonin-related peptide (ACP) gene, previously missing from the draft genome**. The image displays an Integrative Genomics Viewer (IGV) screenshot of the *G. bimaculatus* chromosome-scale assembly. The complete gene model for ACP (Gbim.chrXG0009740.1), one of the nine neuropeptide genes reported missing from the first assembly report (Mochizuki et al., 2023), is shown. The gene is now successfully anchored and annotated on Chromosome X (chrX), spanning a region of approximately 20 kb. The blue track (Gbim_v2.2.gff3) shows the full exon-intron structure (exons as thick blocks, introns as thin lines).

## Notes

### Competing Interest Statement

The authors have declared no competing interest.

