## Supplementary material for "Chromosome-scale genome assembly and annotation of the two-spotted cricket *Gryllus bimaculatus* (Orthoptera: Gryllidae)": Fig. S1, Fig. S2

**Supplementary Figures**

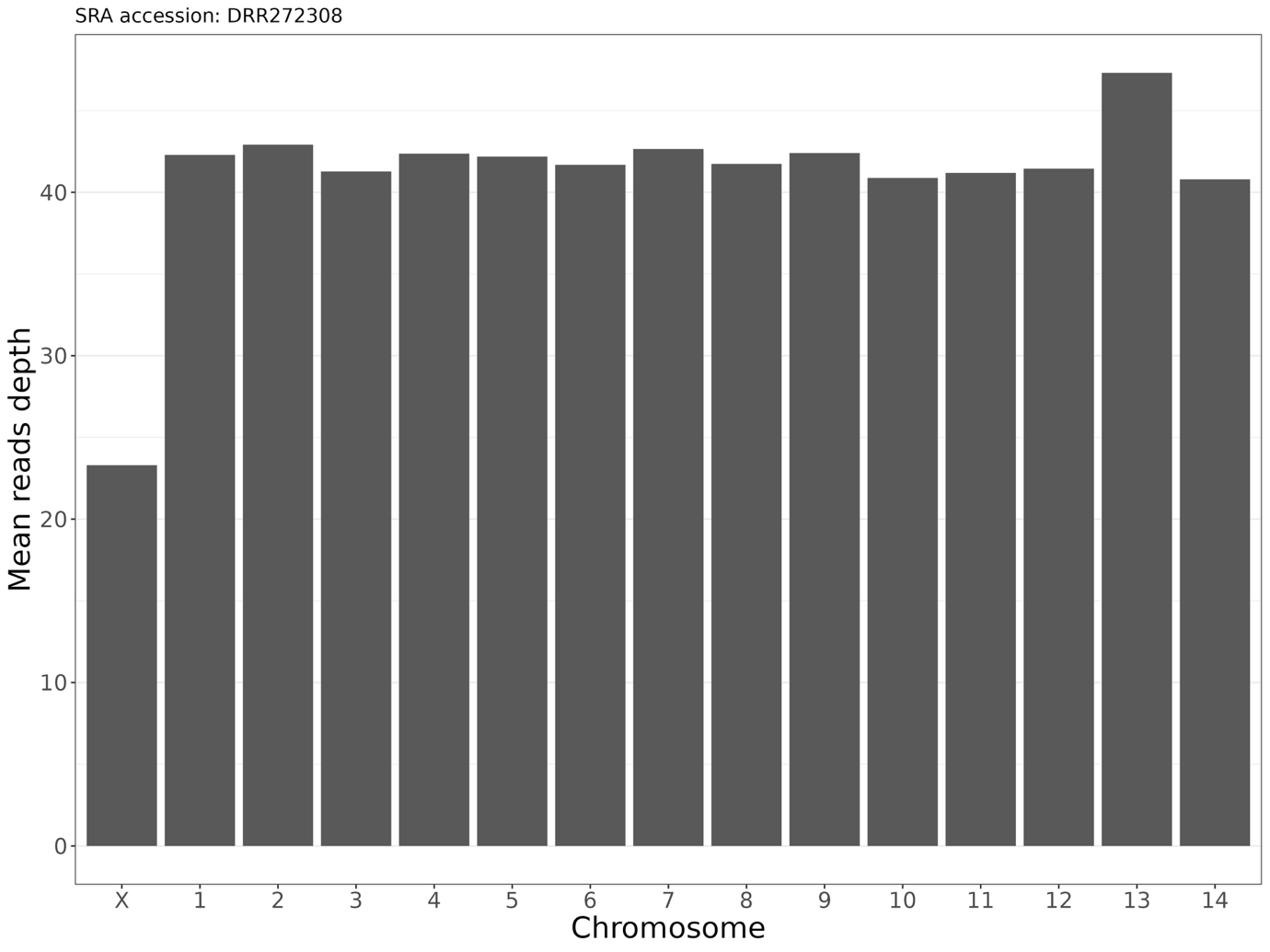

**Supplementary Figure S1. Genomic read coverage confirms X chromosome hemizygosity.**

Mean read coverage depth across all assembled chromosomes derived from the mapping of genomic short reads of a single male *Gryllus bimaculatus* individual (SRA: DRR272308). The X chromosome displays approximately half the average autosomal coverage depth, consistent with the expected pattern of male hemizygosity.

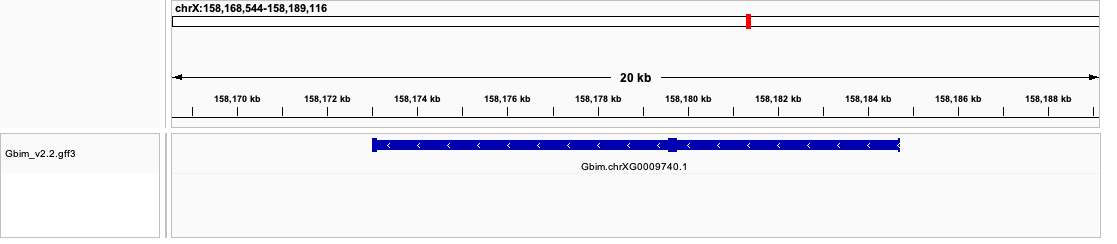

**Supplementary Figure S2. Recovery of the Adipokinetic hormone/corazonin-related peptide (ACP) gene, previously missing from the draft genome.**

The image displays an Integrative Genomics Viewer (IGV) screenshot of the *G. bimaculatus* chromosome-scale assembly. The complete gene model for ACP (Gbim.chrXG0009740.1), one of the nine neuropeptide genes reported missing from the first assembly report (Mochizuki et al., 2023), is shown. The gene is now successfully anchored and annotated on Chromosome X (chrX), spanning a region of approximately 20 kb. The blue track (Gbim_v2.2.gff3) shows the full exon-intron structure (exons as thick blocks, introns as thin lines).
